# FlaHMM: unistrand *flamenco*-like piRNA cluster prediction in *Drosophila* species using hidden Markov models

**DOI:** 10.1101/2024.05.14.592433

**Authors:** Maria-Anna Trapotsi, Jasper van Lopik, Gregory J Hannon, Benjamin Czech Nicholson, Susanne Bornelöv

## Abstract

PIWI-interacting RNAs are a class of small non-coding RNAs that are essential for transposon control in animal gonads. In *Drosophila* ovarian somatic cells, piRNAs are transcribed from large genomic regions called piRNA clusters, which are enriched for transposon fragments and acts as a memory of past invasions. Despite being widely present across *Drosophila* species, somatic piRNA clusters are notoriously difficult to identify and study due to their lack of sequence conservation and limited synteny. Current identification methods rely either on extensive manual curation or availability of high-throughput small RNA-seq data, limiting large-scale comparative studies. We now present FlaHMM, a hidden Markov model developed to automate genomic annotation of *flamenco*-like unistrand piRNA clusters in *Drosophila* species without the need of experimental data beyond a genome assembly. FlaHMM uses transposable element content across 5 or 10 kb bins calculated from genome sequence alone and is thus able to detect candidate piRNA clusters without the need to obtain flies and experimentally perform small RNA sequencing. We show that FlaHMM performs on par with piRNA-guided or manual methods, and thus provides a scalable and efficient approach to piRNA cluster annotation in new genome assemblies. FlaHMM is freely available at https://github.com/Hannon-lab/FlaHMM under an MIT licence.

## Introduction

Transposable elements (TEs) are DNA sequences with the ability to move and amplify within a genome, thus posing a threat to genome integrity of their host. In the fruit fly *D. melanogaster*, the *flamenco* (*flam*) locus plays an essential role in repressing a subset of TEs in somatic follicle cells of the ovary. Here, *flam* serves as the predominant source of PIWI-interacting RNAs (piRNAs), a class of small non-coding RNAs that guide PIWI proteins to silence TEs through complementary base pairing. Failure to express or process *flam* into piRNAs typically results in sterility (Goriaux *et al*., 2014; Prud’homme *et al*., 1995).

Although most animals rely on the piRNA pathway to repress TEs, *flam* was described as a master regulator of *Gypsy*-family TEs in *Drosophila* (Prud’homme *et al*., 1995; Kim *et al*., 1994) a decade before it was known to be a piRNA cluster. For a long time, *flam*-syntenic clusters were identified only in species closely related to *D. melanogaster* (Chirn *et al*., 2015; Malone *et al*., 2009). We recently reported that *flam* is evolutionarily conserved beyond the *melanogaster* subgroup and that *flam*-like loci exist in species that diverged from *D. melanogaster* at least 33 million years ago (van Lopik *et al*., 2023). This raises the possibility that unistrand piRNA clusters may control *Gypsy*-family TEs across the whole *Drosophila* genus.

Currently, piRNA clusters are typically identified by mapping piRNAs onto the genome of interest, followed by identification of candidate piRNA clusters using proTRAC (Rosenkranz and Zischler, 2012). proTRAC quantifies small RNA abundance per 1 kb genomic bin and identifies regions where small RNAs display piRNA-like characteristics such as 1U and 10A biases. Although powerful, proTRAC and related methods crucially relies on the availability of small RNA-seq data obtained from germline cells of the species of interest. Recent advances in long-read sequencing have enabled more and better genome assemblies. High-quality assemblies are now available for 298 Drosophilid species (Kim *et al*., 2023), including many assemblies from individual wild-caught flies. Using small RNA sequencing to identify piRNA clusters is therefore not possible, both due to the sheer scale of the project, and because many of these species are not currently available and may not thrive in laboratory conditions. To effectively study the evolution of piRNA clusters across all Drosophilids, we would therefore need automated methods capable of annotating *flam*-like piRNA clusters genome-wide from genome sequence alone.

Here we present FlaHMM, a hidden Markov model (HMM) that accurately predicts the location of *flam*-like unistrand piRNA clusters solely based on genomic sequence and predicted TE annotations. Inspired by other genome-wide annotation tools such as ChromHMM (Ernst and Kellis, 2012), FlaHMM divides each chromosome into a series of consecutive bins. Each bin is considered by have a hidden state (piRNA cluster or not) giving rise to an observable feature, in this case TE content. The classification task is formulated as deriving the hidden states, based on the observed TE content. HMMs are particularly suited for this task since they correctly assume that the state of each bin simultaneously depends on the previous and on the next bin. FlaHMM was trained using *flam* and *flam*-syntenic regions from six species in the *melanogaster* subgroup, and evaluated on four additional species from the *melanogaster* subgroup and 12 distantly related species with evolutionarily distinct *flam*-like piRNA clusters. Overall, FlaHMM achieved a true positive rate (TPR) of 0.80±0.34 and false positive rate (FPR) of 0.011±0.011, and the predicted clusters showed strong agreement with high-throughput profiling of soma-enriched ovarian piRNAs.

## Methods

### Data and annotations

#### Training and test sets

All genome assemblies used for training or model evaluation are listed in Table S1. In short, the model was trained on species from the *melanogaster* subgroup. Since training required information about the chromosome arms, we used six species with this information available as a training set and the remaining four species as a test set. Another 12 more distantly related species with non-syntenic clusters were used as a second fully independent test set.

#### Evaluation of genome assembly quality

To estimate the quality of each genome assembly, we used the NX metric, which is defined as the length of the shortest contig for which longer and equal length contigs cover at least X% of the assembly (see “01_assembly_stats” in the supplement repository).

#### De novo transposon annotations using EDTA

De novo transposon libraries were constructed using EDTA (v1.9.3) as previously described (van Lopik *et al*., 2023). The genome was divided into 2.5 kilobases (kb), 5 kb, or 10 kb bins using “bedtools makewindows”. *Gypsy*-family transposons were separated by strand and coverage per bin was quantified using “bedtools coverage” and normalised to bin size as previously described (van Lopik *et al*., 2023). For detailed instructions, see “examples” in the FlaHMM repository.

### Hidden Markov model

#### States

Each genomic bin was assigned one of three possible states: none (0), *flam*-like cluster (1), and centromere-like region (2). As ground truth we used previously reported cluster coordinates (van Lopik *et al*., 2023). Centromeric regions were defined at chr2 and chr3 as described in Supplementary Methods S1. Any other bin was considered to be ‘none’.

#### Emissions

Each genomic bin was assigned one of three possible emissions: no *Gypsy*-family TEs (0), *Gypsy*-family TEs present on one strand (1), *Gypsy*-family TEs on both strands (2). Assignment was done based on the EDTA-predicted *Gypsy*-family TE coverage per strand, defined as a fraction between 0 and 1. This fraction was compared to a threshold between 0.025 and 0.900 using the conditions described in Table S2.

#### Model training

Transition, emission and starting probabilities were estimated as maximum likelihood estimates using the known states and emissions for *D. mauritiana, D. melanogaster, D. santomea, D. simulans, D. subpulchrella*, and *D. yakuba* with pseudo-count 0.001 for transition probabilities, or 1 for emission probabilities and starting probabilities. Finally, a combined model was constructed as the mean of the individual models. Examples of transition matrix (Figure 1A), starting (Figure 1B) and emission matrix (Figure S2) probabilities are provided.

**Figure 1:**
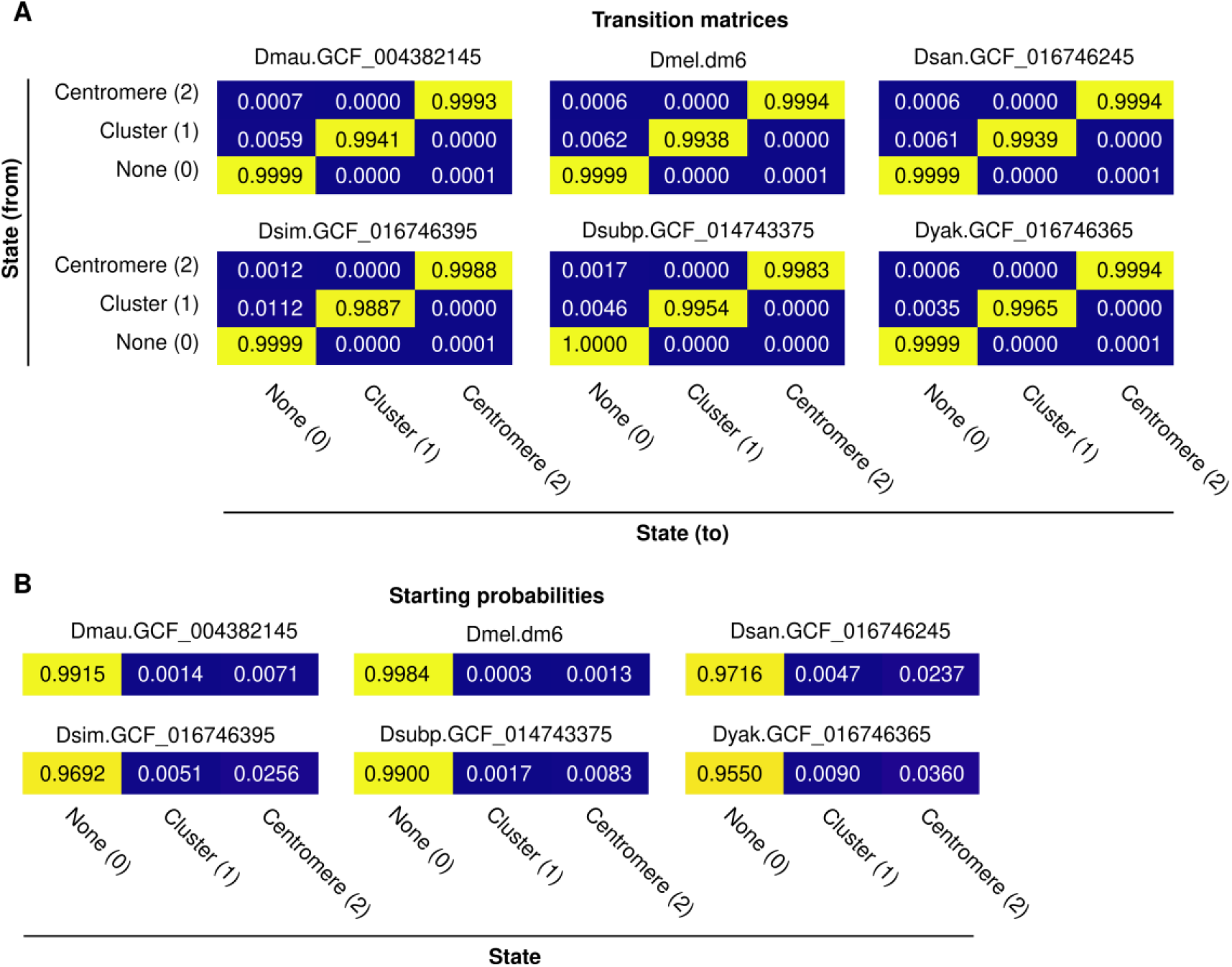
Overview of model parameters. (A) Transition matrix probabilities and (B) starting probabilities calculated across all six *Drosophila* species used to train the models, using the 5 kb binning strategy. Please note that the state is independent of the emission threshold.

#### Model evaluation

The models were implemented in Python using CategoricalHMM (called MultinomialHMM prior to hmmlearn v0.2.8) from hmmlearn (v0.2.7). Parameter optimization during training was done by leave-one-out cross validation. The final models were evaluated on an external test set that included four species with a *flam*-syntenic cluster and 12 species with a previously predicted *flam*-like locus that lacks synteny to *flam*. Models were evaluated by TPR, FPR, precision (eq. 1), recall (eq. 2) and F1-score (eq. 3).

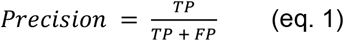

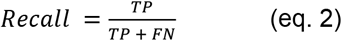

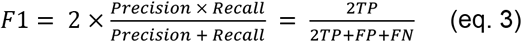

We note that false positives were more common on fragmented and unplaced contigs. We therefore present the cross-validation results across the N90 contigs. See Supplementary Results S1 for more details on NX thresholds.

### Analysis of sRNA-seq data

Sequencing data for *D. ficusphila* available on GEO (accession GSM7059862 and GSM7059863) was processed and aligned as previously described (van Lopik *et al*., 2023). In short, we excluded an abundant rRNA and performed adapter trimming using Trim Galore! (v0.6.4, --stringency 30 -e 0.1 -a TGCTTGGACTACATATGGTTGAGGGTTGTA --length 18 -q 0, followed by --stringency 5 –e 0.1 --length 18 --max_length 35 -q 0). Next, we used bowtie (v1.2.3) to exclude reads mapping to miRBase release 22.1 (Kozomara *et al*., 2019) (-S -n 2 -M 1 -p 20 --best --strata --nomaqround --chunkmbs 1024), followed by aligning the remaining reads to the reference genomes (-S -n 2 -M 1 -p 20 --best --strata --nomaqround --chunkmbs 1024). piRNA cluster prediction was performed using proTRAC (Rosenkranz and Zischler, 2012). Two biological replicates were combined using samtools merge, following by running proTRAC (v2.4.4, -pdens 0.01 -swincr 100 -swsize 1000 -clsize 5000 -1To10A 0.75 -clstrand 0.5 -pimin 23 -pimax 30 -pisize 0.75 -distr 1-99 - nomotif -format SAM).

To quantify the number of piRNAs per genomic region, we further used “bedtools makewindows” with “-w 10000 -s 5000” to construct 10 kb windows with 5 kb overlap. The BAM file was filtered for reads within the expected piRNA size range (24-29 nucleotides), downsampling to at most 1000 reads per 5’ end position. The BAM files were then converted to BED using “bedtools bamtobed” and the number of piRNAs mapping to each bin was quantified using “bedtools intersect” with “-c -F 0.5”.

## Results

### Design of FlaHMM to accurately identify *flam*-like clusters from genomic sequence

To identify the best strategy for *flam*-like cluster identification using *Gypsy*-family TE content (Figure 2A), we trained HMMs using six *Drosophila* species and used leave-one-out cross validation to evaluate how genome binning strategy (2.5, 5, or 10 kb) and *Gypsy*-family TE content thresholds (0.025-0.900) influenced model performance (Figure S4A-C). We note that 5 kb or 10 kb bins gave similar performance (Figure S5B-C), but that lower performance was obtained with 2.5 kb bins (Figure S5A). For all binning strategies, low *Gypsy*-family TE thresholds (0.025-0.100) performed better than higher thresholds (0.200-0.900). For example, the median F_1_-score was equal to 0.79, 0.64 and 0.59 for thresholds 0.025, 0.50 and 0.90, respectively, when 5 kb bins were used (Figure S4B). Based on cross validation F_1_-scores, we selected the top three models using 5 kb bins (thresholds 0.025, 0.5, and 0.075) and the top three models using 10 kb bins (thresholds 0.05, 0.075, and 0.1), resulting in six preferred settings (Figure 2B, and arrows in Figure S4). The overall best performing model (5 kb bins, 0.025 threshold) had median precision 0.76, recall 0.86, and F_1_-score 0.79. To not overestimate performance due to uneven class distribution, all metrics are reported as mean across all three classes (more details in Supplementary Results S2).

**Figure 2:**
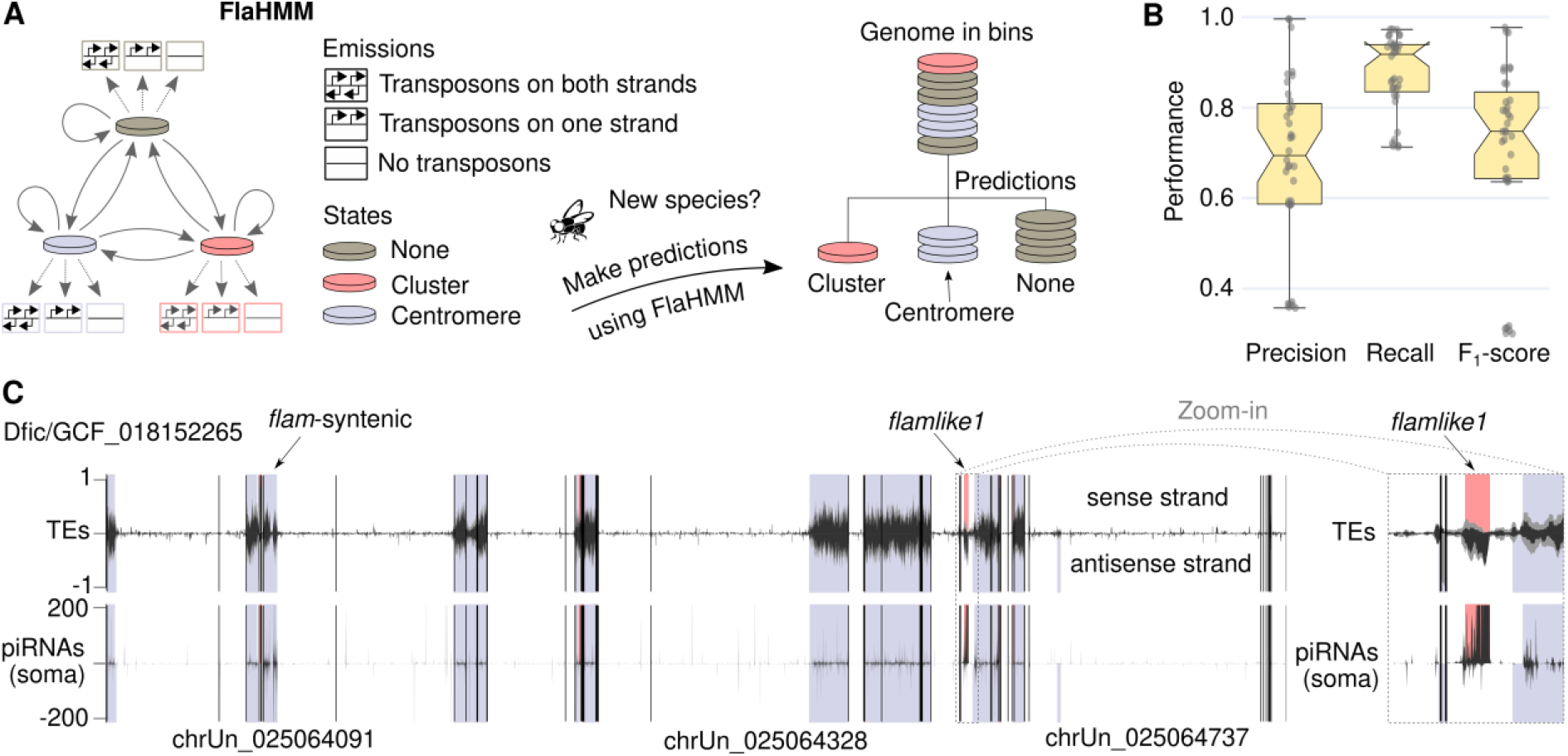
Overview of FlaHMM. (A) Schematic of FlaHMM, an HMM trained to predict unistrand piRNA clusters and centromere-like regions based on *Gypsy*-family TE content per genomic strand. Predictions are made across 2.5-10 kb genomic bins. (B) Cross validation performance across six top-performing models. (C) FlaHMM predictions across the *D. ficusphila* genome. Genomic coordinates are shown on the x axis with vertical black lines indicating contig breaks and selected contig names indicated. The top tracks show transposon content (black lines, LTR TEs; grey lines, all TEs) and the bottom one (black lines) shows piRNAs in soma-enriched ovarian cells (van Lopik *et al*., 2023). Positive values represent the sense strand and negative values the antisense strand relative to *flamlike1*. FlaHMM predictions are shown as shaded areas, with grey-blue regions indicating centromeres, and pink areas *flam*-like clusters. Notably, the only major *flam*-like cluster predicted corresponds to previously reported *flamlike1* (van Lopik *et al*., 2023), indicated by an arrow and shown in the zoom-in to the right (grey dashed lines).

### FlaHMM efficiently scans *Drosophila* genomes for *flam*-like piRNA cluster candidates

FlaHMM output includes interactive plots (Figure S6-S8) that can be explored in a web browser. We next applied FlaHMM on 29 genome assemblies from 16 previously unseen *Drosophila* species with previously reported unistrand piRNA clusters. Four of these species belong to the *melanogaster* species subgroup and have *flam*-syntenic piRNA clusters, and another 12 are evolutionarily distant with non-syntenic *flam*-like clusters (van Lopik *et al*., 2023).

We first used the settings that performed best during cross validation (5 kb bins, 0.025 threshold). Known *flam*-syntenic clusters were successfully re-identified in 11 out of 11 assemblies representing all four tested species (Table S3, overall TPR 0.900±0.111, FPR 0.011±0.013). For the more difficult task of finding new clusters, we re-identified *flam*-like clusters in 14 out of 18 assemblies from nine species (Table S4, overall TPR 0.737±0.411, FPR 0.010±0.011), representing four distinct unistrand piRNA clusters (*flamlike1, flamlike2, flamlike3*, and *flamlike5*). We note that although even a low FPR can pose a problem when predicting a rare category on a genome-wide scale, these were generally scattered across the centromere-like regions and therefore easy to separate from the pericentromeric piRNA cluster (Figure 2C). Notably, our successful predictions included *D. ambigua, D. miranda, D. obscura*, and *D. tristis*, where previously reported *flam*-like regions were identified only when guided by synteny analysis (van Lopik *et al*., 2023). FlaHMM therefore provides an improvement in sensitivity over the currently used manual annotation.

We next evaluated FlaHMM using the other five top-performing settings (Table 1). Strikingly, while all settings re-identified 11 out of 11 *flam*-syntenic regions (Table S5), the new settings, which all used higher thresholds, performed equally well or better on non-syntenic *flam*-like clusters. For instance, using 5 kb bins and 0.075 threshold, we successfully identified *flam*-like clusters in 28 out of 29 genome assemblies (Table S5 and Table S6, overall TPR 0.898±0.195, FPR 0.014±0.017), failing only on the highly fragmented Dfic.GCF_000220665 assembly (5,754 contigs and N50 of 1,050,541). All metrics are shown in Table S7. We conclude that higher thresholds likely improve identification of non-syntenic *flam*-like clusters, although this remains to be verified on independent data. We nevertheless recommend running FlaHMM with higher thresholds if no hit is found initially.

**Table 1:** Model performance overview. Results shown for N90 contigs, except for Damb.d101g, which uses N100. Detailed metrics are available in Table S7.

| Model | Bin | Threshold | TPR | FPR |
| --- | --- | --- | --- | --- |
| 5kb_CV_rank1 | 5 kb | 0.025 | 0.799 ± 0.336 | 0.011 ± 0.011 |
| 5kb_CV_rank2 | 5 kb | 0.05 | 0.795 ± 0.324 | 0.010 ± 0.011 |
| 5kb_CV_rank3 | 5 kb | 0.075 | 0.898 ± 0.195 | 0.014 ± 0.017 |
| 10kb_CV_rank1 | 10 kb | 0.05 | 0.832 ± 0.269 | 0.019 ± 0.019 |
| 10kb_CV_rank2 | 10 kb | 0.1 | 0.868 ± 0.218 | 0.019 ± 0.021 |
| 10kb_CV_rank3 | 10 kb | 0.075 | 0.833 ± 0.267 | 0.018 ± 0.019 |

### FlaHMM predictions are experimentally supported

Next, we tested the agreement between FlaHMM predictions and piRNA profiling through small RNA sequencing. In *D. ficusphila*, FlaHMM identified 91% of bins overlapping *flamlike1*, limiting FPR to 0.5% located primarily in centromeric regions (Figure 2C). Publicly available soma-enriched piRNA profiling (van Lopik *et al*., 2023) confirmed that this region was strongly enriched for piRNAs originating from the predicted strand and complementary to *Gypsy*-family TEs, thus resembling the expression pattern of a unistrand piRNA cluster. In comparison, using the same sequencing data, proTRAC predicted 50 piRNA clusters (Table S8), including 13 clusters overlapping *flamlike1*, and recovered 50% of bins overlapping *flamlike1* at a 0.3% FPR (Figure S9). Thus, both approaches are largely successful, although the latter requires experimental sequencing data.

## Conclusion

We have developed FlaHMM, a tool for unistrand piRNA cluster prediction in *Drosophila* species. Notably, while piRNA cluster identification traditionally has relied on small RNA sequencing, FlaHMM predicts cluster locations with no experimental data beyond a genome assembly. We hope that the release of FlaHMM will simplify and speed up annotation of unistrand piRNA cluster candidates in newly assembled *Drosophila* genomes. Moreover, with FlaHMM as a proof-of-principle, similar tools may now be developed for other species.

## Code Availability

We have released two GitHub repositories. The main repository contains the FlaHMM tool and instructions on how to run it (https://github.com/Hannon-lab/FlaHMM) (M.-A. Trapotsi *et al*., 2024a). The second repository contains all the supplementary information figures and scripts used to create them (https://github.com/Hannon-lab/FlaHMM-supplement) (M.-A. Trapotsi *et al*., 2024b).

## Data Availability

All data used are publicly available. Sequencing data for *D. ficusphila* is available on GEO (accession GSM7059862 and GSM7059863). Genome assemblies referred to as ‘d15genomes’ (Miller *et al*., 2018) or ‘d101g’ (Kim *et al*., 2021) can be downloaded from their original publication, all other assemblies are referred to by their accession number (GCA for GenBank assemblies and GCF for RefSeq).

## Acknowledgements

We thank members of the Hannon and Bornelöv labs for fruitful discussions, and the Scientific Computing core facility at CRUK Cambridge Institute for HPC resources.

## Author Contributions

SB conceived the study and supervised the study together with BCN and GJH. MAT implemented the tool, prepared Jupyter notebooks and performed statistical analyses. SB collected and processed raw data and assisted with software development and documentation. JvL validated the final code. MAT and SB drafted the manuscript with input from all authors. All authors interpreted the results and approved the final manuscript.

