## Supplementary Information for "FlaHMM: unistrand *flamenco*-like piRNA cluster prediction in *Drosophila* species using hidden Markov models"


**Supplementary Methods**
**Methods S1**: Centromere definition.

**Supplementary Results**
**Results S1:** Model performance depends on contig size.
**Results S2**: Model evaluation and class imbalance.

**Supplementary Tables**
**Table S1:** Overview of species included in the training and test sets.
**Table S2:** Definition of emissions.
**Table S3:** Performance on genome assemblies with *flam*-syntenic clusters.
**Table S4:** Performance on genome assemblies with *flam*-like clusters.
**Table S5:** Performance on genome assemblies with *flam*-syntenic clusters [5 kb bins, 0.075 threshold settings].
**Table S6:** Performance on genome assemblies with *flam*-like clusters [5 kb bins, 0.075 threshold settings].
**Table S7:** Detailed performance for top six model settings.
**Table S8:** proTRAC piRNA cluster predictions for *D. ficusphila*.

**Supplementary Figures**
**Figure S1:** Automatic annotation of centromeric regions.
**Figure S2:** Emission matrices depend on TE content threshold.
**Figure S3:** Overview of model performance by NX threshold and bin size.
**Figure S4:** Overview of model performance by threshold and bin size.
**Figure S5:** Additional details on model performance.
**Figure S6:** Model output for *D. oshimai*.
**Figure S7:** Model output for *D. innubila*.
**Figure S8:** Model output for *D. ficusphila*.
**Figure S9:** Overview of proTRAC predictions for *D. ficusphila*.

**Supplementary References**

**Methods S1: Centromere definition**

Centromeres in *D. melanogaster* are enriched for non-LTR retroelements belonging to the Jockey family, particularly G2/Jockey-3, although these elements are not exclusive to centromeres(Chang *et al.*, 2019). Similar enrichment is observed in other species, such as *D. simulans*, where enrichment of G2/Jockey-3 elements at centromeres has been observed (Talbert *et al.*, 2018). This suggests a conserved role for retroelements in centromeric function, as proposed for other species (Klein and O'Neill, 2018).

While most transposon families, including LTR elements, are present in centromeres, distinguishing these regions from *flam*-like clusters is crucial. To achieve this, we annotated centromeric regions separately, allowing FlaHMM to discern them from *flam*-like regions. Genome assemblies with contigs labelled as the left and right arm of a chromosome (i.e., chr2L and chr2R, or chr3L and chr3R) were used in combination with predicted TE content to define centromeric regions. Each species' genome was binned into 100 kb bins and TE content was calculated per bin. A sliding window of 5 bins was used to evaluate TE content, with centromeric regions defined to start at the chromosome end and extend for as long as the mean TE content across the 5 bins was exceeding 20%. A visualisation of this calculation for *D. melanogaster* is shown in Figure S1, with metrics for all species available in the GitHub repository (centromere_definition, https://github.com/Hannon-lab/FlaHMM-supplement).

**Results S1: Model performance depends on contig size**

To systematically determine how performance depends on contig size, we evaluated the model on a subset of the full assemblies using N80, N85, N90, N95, N99, and N100 as contig cutoff (Figure S3). Notably, performance substantially improved when limiting evaluation to contigs corresponding to the 90th or higher most continuous percentile of the assembly, compared to using the full assembly (N100). We therefore concluded that most prediction errors occur at small and unplaced contigs, and although predictions can be made for all contigs, we recommend the exclusion of these when interpreting the results.

**Results S2: Model evaluation and class imbalance**

We note that representing and interpreting classification performance is challenging in situations with uneven class distribution, such as the one presented in our study. Since the class distribution was highly uneven (e.g. 87.6% None, 0.3% Cluster and 12.1% Centromere for *D. melanogaster*), we therefore represented model performance as the mean performance across all three classes throughout our study. However, it should be noted that performance varies substantially by class (Figure S5C), with larger classes obtaining higher performance, and that the global performance (weighted) is substantially better than the mean performance (Figure S5C).

**Table S1: Overview of species included in the training and test sets.** Annotated unistrand piRNA cluster coordinates were used as previously reported (van Lopik *et al.*, 2023). Test refers to *flam*-syntenic clusters, whereas Test2 refers to *flam*-like ones without synteny.

| | | | **Unistrand piRNA cluster** | | | | |
|---|---|---|---|---|---|---|---|
| **Dataset** | **Species** | **Assembly** | **Name** | **Contig** | **Start** | **End** | **Strand** |
| Training | Dyak | GCF_016746365 | *flam* | chrX | 21,210,985 | 22,624,858 | + |
| Training | Dsan | GCF_016746245 | *flam* | chrX | 21,033,264 | 21,845,911 | + |
| Training | Dsim | GCF_016746395 | *flam* | chrX | 21,073,454 | 21,511,484 | + |
| Training | Dmau | GCF_004382145 | *flam* | chrX | 21,407,318 | 22,252,287 | + |
| Training | Dmel | dm6 | *flam* | chrX | 21,631,904 | 22,434,871 | + |
| Training | Dsubp | GCF_014743375 | *flam* | chrX | 1,890,097 | 2,975,514 | - |
| Test | Dbia | GCF_018148935 | *flam*-syntenic | chrUn_025319364 | 24,741,581 | 24,969,825 | + |
| Test | Dbia | d101g | *flam*-syntenic | contig_275 | 24,743,392 | 24,969,643 | + |
| Test | Dbia | d15genomes | *flam*-syntenic | utg000088l | 779,356 | 972,521 | - |
| Test | Dere | GCF_003286155 | *flam*-syntenic | chrUn_020825209 | 15,529,294 | 15,801,676 | + |
| Test | Dere | d101g | *flam*-syntenic | contig_636 | 419,529 | 994,684 | - |
| Test | Dere | droEre1 | *flam*-syntenic | scaffold_4690 | 17,965,809 | 18,220,474 | + |
| Test | Dere | d15genomes | *flam*-syntenic | utg000005l | 2,667,515 | 2,942,695 | + |
| Test | Dsuz | GCF_013340165 | *flam*-syntenic | chrX_023496844 | 665,099 | 1,345,189 | + |
| Test | Dtei | GCF_016746235 | *flam*-syntenic | chrX | 20,998,347 | 21,767,495 | + |
| Test | Dtei | d101g_2733 | *flam*-syntenic | contig_529 | 1 | 180,271 | - |
| Test | Dtei | d101g_CT02 | *flam*-syntenic | contig_903 | 563,752 | 778,755 | - |
| Test2 | Dfic | GCF_018152265 | *flamlike1* | chrUn_025064569 | 438,589 | 994,838 | - |
| Test2 | Dfic | d101g | *flamlike1* | contig_636 | 419,529 | 994,684 | - |
| Test2 | Dfic | GCF_000220665 | *flamlike1* | chrUn_016073254 | 1 | 77,405 | - |
| Test2 | Dosh | d101g | *flamlike2* | contig_1 | 23,542,221 | 24,075,495 | + |
| Test2 | Dath | GCA_008121215 | *flamlike3* | chrX | 35,768,372 | 36,552,428 | + |
| Test2 | Dazt | GCA_005876895 | *flamlike3* | VCKU01000012 | 4,140,196 | 4,501,485 | + |
| Test2 | Dmir | GCF_003369915 | *flamlike3* | chrXR | 30,822,007 | 31,271,755 | - |
| Test2 | Dper | GCF_003286085 | *flamlike3* | chrUn_020825317 | 2,171,578 | 2,418,435 | + |
| Test2 | Dper | d101g | *flamlike3* | contig_44 | 385,239 | 505,116 | + |
| Test2 | Dper | d15genomes | *flamlike3* | utg000136l | 407,796 | 438,716 | + |
| Test2 | Dpse | d15genomes | *flamlike3* | utg000003l | 435,828 | 675,327 | + |
| Test2 | Dpse | GCF_009870125 | *flamlike3* | chrX | 37,962,257 | 38,275,576 | + |
| Test2 | Dinn | GCF_004354385 | *flamlike4* | chrX | 30,916,784 | 31,158,180 | - |
| Test2 | Damb | d101g | *flamlike5* | contig_109 | 126,545 | 295,508 | - |
| Test2 | Dbif | GCA_009664405 | *flamlike5* | chrA | 13,681,722 | 13,800,479 | - |
| Test2 | Dobs | d101g | *flamlike5* | contig_185 | 1 | 66,608 | - |
| Test2 | Dobs | GCF_018151105 | *flamlike5* | chrUn_024542773 | 1 | 56,444 | - |
| Test2 | Dtris | d101g | *flamlike5* | contig_125 | 2,204,925 | 2,260,492 | + |

**Table S2: Definition of emissions**. Emissions were defined based on three conditions relative to a threshold (*τ*) as specified below. ***G+*** and ***G-*** represent the Gypsy-family TE content (G) on the plus (+) and minus (-) strand, respectively, represented as a fraction of the total bin size.

| LTR/Gypsy content | | | Emission | |
|---|---|---|---|---|
| ***G+*** − ***G-*** > *τ* | ***G-*** − ***G+*** > *τ* | ***G+*** + ***G-*** > *τ* | Plus strand | Minus strand |
| FALSE | FALSE | FALSE | 0 (none) | 0 (none) |
| FALSE | FALSE | TRUE | 2 (both) | 2 (both) |
| FALSE | TRUE | | 1 (one) | 0 (none) |
| TRUE | FALSE | | 0 (none) | 1 (one) |

**Table S3: Performance on genome assemblies with *flam*-syntenic clusters.** Performance for the 5 kb bin, 0.025 threshold settings is shown as TPR and FPR, respectively. Species are sorted alphabetically and multiple assemblies for the same species are sorted based on N50 as an indication of assembly quality. Results shown for N90 contigs unless otherwise specified. Detailed metrics are shown in Table S7.

| **Assembly** | **Cluster** | **TPR** | **FPR** | **Contigs** | **N50** | **Total size** |
|---:|---:|---:|---:|---:|---:|---:|
| Dbia.GCF_018148935 | *flam*-syntenic | 1.0000 | 0.0177 | 283 | 23,381,765 | 185,318,521 |
| Dbia.d101g | *flam*-syntenic | 1.0000 | 0.0179 | 283 | 23,381,765 | 185,318,521 |
| Dbia.d15genomes | *flam*-syntenic | 0.9750 | 0.0143 | 661 | 2,791,184 | 182,453,935 |
| Dere.GCF_003286155 | *flam*-syntenic | 0.7500 | 0.0032 | 94 | 22,146,549 | 146,538,397 |
| Dere.d101g | *flam*-syntenic | 0.9211 | 0.0002 | 499 | 20,389,455 | 135,657,613 |
| Dere.droEre1 | *flam*-syntenic | 0.7692 | 0.0004 | 5,124 | 18,750,251 | 152,862,534 |
| Dere.d15genomes | *flam*-syntenic | 0.7500 | 0.0000 | 58 | 16,960,765 | 130,293,209 |
| Dsuz.GCF_013340165 | *flam*-syntenic | 1.0000 | 0.0411 | 546 | 2,609,782 | 268,012,156 |
| Dtei.GCF_016746235 | *flam*-syntenic | 0.7806 | 0.0000 | 91 | 24,551,875 | 149,494,471 |
| Dtei.d101g_2733 | *flam*-syntenic | 0.9730 | 0.0186 | 831 | 8,218,613 | 152,423,477 |
| Dtei.d101g_CT02 | *flam*-syntenic | 0.9773 | 0.0102 | 1,144 | 7,096,444 | 144,692,533 |

**Table S4: Performance on genome assemblies with *flam*-like clusters.** Performance for the 5 kb bin, 0.025 threshold settings is shown as TPR and FPR, respectively. Species are sorted alphabetically and multiple assemblies for the same species are sorted based on N50 as an indication of assembly quality. Results shown for N90 contigs unless otherwise specified. Detailed metrics are shown in Table S7.

| Assembly | Cluster | TPR | FPR | Contigs | N50 | Total size |
|---:|---:|---:|---:|---:|---:|---:|
| Dfic.GCF_018152265 | *flamlike1* | 0.9196 | 0.0052 | 838 | 9,958,166 | 167,832,931 |
| Dfic.d101g | *flamlike1* | 0.8879 | 0.0049 | 838 | 9,958,166 | 167,832,931 |
| Dfic.GCF_000220665 | *flamlike1* | 0.0000 | 0.0024 | 5,754 | 1,050,541 | 152,439,475 |
| Dosh.d101g | *flamlike2* | 1.0000 | 0.0034 | 323 | 2,625,980 | 181,013,079 |
| Dath.GCA_008121215 | *flamlike3* | 0.0000 | 0.0017 | 119 | 52,101,127 | 192,660,667 |
| Dazt.GCA_005876895 | *flamlike3* | 0.0000 | 0.0102 | 126 | 17,833,963 | 219,083,522 |
| Dmir.GCF_003369915 | *flamlike3* | 1.0000 | 0.0329 | 104 | 35,263,383 | 287,096,000 |
| Dper.GCF_003286085 | *flamlike3* | 1.0000 | 0.0362 | 432 | 5,212,974 | 195,512,972 |
| Dper.d101g | *flamlike3* | 1.0000 | 0.0073 | 1,072 | 4,059,666 | 152,810,519 |
| Dper.d15genomes | *flamlike3* | 1.0000 | 0.0166 | 415 | 3,429,058 | 163,933,157 |
| Dpse.d15genomes | *flamlike3* | 0.9796 | 0.0225 | 361 | 2,983,193 | 159,031,139 |
| Dpse.GCF_009870125 | *flamlike3* | 0.8906 | 0.0198 | 70 | 32,422,566 | 163,282,969 |
| Dinn.GCF_004354385 | *flamlike4* | 0.0000 | 0.0003 | 363 | 29,570,363 | 167,978,031 |
| Damb.d101g | *flamlike5* | 0.9714[1] | 0.0053[1] | 172 | 11,334,561 | 161,264,264 |
| Dbif.GCA_009664405 | *flamlike5* | 0.7600 | 0.0040 | 214 | 48,071,510 | 192,749,618 |
| Dobs.d101g | *flamlike5* | 0.8571 | 0.0121 | 215 | 3,931,141 | 179,834,232 |
| Dobs.GCF_018151105 | *flamlike5* | 1.0000 | 0.0079 | 215 | 3,931,141 | 179,834,232 |
| Dtris.d101g | *flamlike5* | 1.0000 | 0.0003 | 275 | 5,059,044 | 158,167,647 |

[1] Results shown for N100 contigs since *flamlike5* is located on a contig below the N90 threshold.

**Table S5: Performance on genome assemblies with *flam*-syntenic clusters [5 kb bins, 0.075 threshold settings].** Performance for the 5 kb bin, 0.075 threshold settings is shown as TPR and FPR, respectively. Species are sorted alphabetically and multiple assemblies for the same species are sorted based on N50 as an indication of assembly quality. Results shown for N90 contigs unless otherwise specified. Detailed metrics are shown in Table S7.

| Assembly | Cluster | TPR | FPR | Contigs | N50 | Total size |
|---:|---:|---:|---:|---:|---:|---:|
| Dbia.GCF_018148935 | *flam*-syntenic | 0.9783 | 0.0116 | 283 | 23,381,765 | 185,318,521 |
| Dbia.d101g | *flam*-syntenic | 0.9783 | 0.0127 | 283 | 23,381,765 | 185,318,521 |
| Dbia.d15genomes | *flam*-syntenic | 0.9500 | 0.0093 | 661 | 2,791,184 | 182,453,935 |
| Dere.GCF_003286155 | *flam*-syntenic | 0.7500 | 0.0067 | 94 | 22,146,549 | 146,538,397 |
| Dere.d101g | *flam*-syntenic | 0.9211 | 0.0002 | 499 | 20,389,455 | 135,657,613 |
| Dere.droEre1 | *flam*-syntenic | 0.6731 | 0.0008 | 5,124 | 18,750,251 | 152,862,534 |
| Dere.d15genomes | *flam*-syntenic | 0.7500 | 0.0005 | 58 | 16,960,765 | 130,293,209 |
| Dsuz.GCF_013340165 | *flam*-syntenic | 1.0000 | 0.0449 | 546 | 2,609,782 | 268,012,156 |
| Dtei.GCF_016746235 | *flam*-syntenic | 1.0000 | 0.0000 | 91 | 24,551,875 | 149,494,471 |
| Dtei.d101g_2733 | *flam*-syntenic | 0.9730 | 0.0189 | 831 | 8,218,613 | 152,423,477 |
| Dtei.d101g_CT02 | *flam*-syntenic | 0.9545 | 0.0077 | 1,144 | 7,096,444 | 144,692,533 |

**Table S6: Performance on genome assemblies with *flam*-like clusters [5 kb bins, 0.075 threshold settings].** Performance for the 5 kb bin, 0.075 threshold settings is shown as TPR and FPR, respectively. Species are sorted alphabetically and multiple assemblies for the same species are sorted based on N50 as an indication of assembly quality. Results shown for N90 contigs unless otherwise specified. Detailed metrics are shown in Table S7.

| Assembly | Cluster | TPR | FPR | Contigs | N50 | Total size |
|---:|---:|---:|---:|---:|---:|---:|
| Dfic.GCF_018152265 | *flamlike1* | 0.9018 | 0.0084 | 838 | 9,958,166 | 167,832,931 |
| Dfic.d101g | *flamlike1* | 0.8707 | 0.0084 | 838 | 9,958,166 | 167,832,931 |
| Dfic.GCF_000220665 | *flamlike1* | 0.0000 | 0.0018 | 5,754 | 1,050,541 | 152,439,475 |
| Dosh.d101g | *flamlike2* | 1.0000 | 0.0040 | 323 | 2,625,980 | 181,013,079 |
| Dath.GCA_008121215 | *flamlike3* | 0.9937 | 0.0020 | 119 | 52,101,127 | 192,660,667 |
| Dazt.GCA_005876895 | *flamlike3* | 1.0000 | 0.0360 | 126 | 17,833,963 | 219,083,522 |
| Dmir.GCF_003369915 | *flamlike3* | 0.9560 | 0.0715 | 104 | 35,263,383 | 287,096,000 |
| Dper.GCF_003286085 | *flamlike3* | 1.0000 | 0.0433 | 432 | 5,212,974 | 195,512,972 |
| Dper.d101g | *flamlike3* | 1.0000 | 0.0093 | 1,072 | 4,059,666 | 152,810,519 |
| Dper.d15genomes | *flamlike3* | 1.0000 | 0.0202 | 415 | 3,429,058 | 163,933,157 |
| Dpse.d15genomes | *flamlike3* | 0.9796 | 0.0233 | 361 | 2,983,193 | 159,031,139 |
| Dpse.GCF_009870125 | *flamlike3* | 0.8906 | 0.0427 | 70 | 32,422,566 | 163,282,969 |
| Dinn.GCF_004354385 | *flamlike4* | 1.0000 | 0.0018 | 363 | 29,570,363 | 167,978,031 |
| Damb.d101g | *flamlike5* | 0.9714[1] | 0.0116[1] | 172 | 11,334,561 | 161,264,264 |
| Dbif.GCA_009664405 | *flamlike5* | 0.7600 | 0.0036 | 214 | 48,071,510 | 192,749,618 |
| Dobs.d101g | *flamlike5* | 0.8571 | 0.0100 | 215 | 3,931,141 | 179,834,232 |
| Dobs.GCF_018151105 | *flamlike5* | 1.0000 | 0.0129 | 215 | 3,931,141 | 179,834,232 |
| Dtris.d101g | *flamlike5* | 0.9231 | 0.0005 | 275 | 5,059,044 | 158,167,647 |

[1] Results shown for N100 contigs since *flamlike5* is located on a contig below the N90 threshold.

**Table S7: Detailed performance for top six model settings.**

- ⇨ Table available in a separate Excel sheet.

**Table S8: proTRAC piRNA cluster predictions for *D. ficusphila***. Clusters overlapping *flamlike1* are shown with an orange background.

| Chromosome | Start | End | Strand | Reads | 1U fraction | 10A fraction | piRNA-sized fraction |
|---|---|---|---|---|---|---|---|
| chrUn_025063985 | 37114 | 44001 | + | 15484 | 0.78 | 0.28 | 0.92 |
| chrUn_025063993 | 449731 | 455926 | - | 11878 | 0.85 | 0.19 | 0.93 |
| chrUn_025064091 | 364257 | 369917 | + | 11198 | 0.86 | 0.35 | 0.92 |
| chrUn_025064091 | 376862 | 382304 | + | 10806 | 0.85 | 0.36 | 0.93 |
| chrUn_025064091 | 1301599 | 1308329 | . | 14939 | 0.86 | 0.37 | 0.93 |
| chrUn_025064091 | 1333411 | 1345128 | + | 32080 | 0.85 | 0.30 | 0.90 |
| chrUn_025064091 | 1420415 | 1429405 | - | 15942 | 0.85 | 0.26 | 0.91 |
| chrUn_025064091 | 1659300 | 1665116 | - | 19783 | 0.75 | 0.16 | 0.94 |
| chrUn_025064091 | 1729905 | 1737103 | + | 17625 | 0.83 | 0.28 | 0.93 |
| chrUn_025064091 | 1793131 | 1799557 | + | 8639 | 0.85 | 0.16 | 0.90 |
| chrUn_025064092 | 1314113 | 1319927 | - | 8224 | 0.87 | 0.33 | 0.95 |
| chrUn_025064092 | 9849861 | 9856888 | + | 19588 | 0.78 | 0.38 | 0.91 |
| chrUn_025064092 | 13781325 | 13789616 | - | 28620 | 0.83 | 0.26 | 0.92 |
| chrUn_025064092 | 13969206 | 13975554 | - | 15898 | 0.84 | 0.15 | 0.93 |
| chrUn_025064092 | 14948810 | 14954280 | . | 7352 | 0.84 | 0.27 | 0.94 |
| chrUn_025064094 | 1457100 | 1466214 | + | 15920 | 0.82 | 0.19 | 0.93 |
| chrUn_025064118 | 480411 | 495894 | + | 32320 | 0.83 | 0.35 | 0.93 |
| chrUn_025064156 | 1099601 | 1104901 | - | 7771 | 0.92 | 0.26 | 0.93 |
| chrUn_025064203 | 330004 | 337808 | + | 19705 | 0.81 | 0.37 | 0.93 |
| chrUn_025064220 | 20507 | 27607 | - | 17042 | 0.84 | 0.13 | 0.95 |
| chrUn_025064220 | 47507 | 54222 | + | 10114 | 0.85 | 0.46 | 0.92 |
| chrUn_025064328 | 13150806 | 13155969 | - | 12001 | 0.81 | 0.36 | 0.90 |
| chrUn_025064328 | 20922251 | 20927720 | + | 14071 | 0.86 | 0.17 | 0.94 |
| chrUn_025064328 | 21103451 | 21110345 | - | 12608 | 0.87 | 0.23 | 0.93 |
| chrUn_025064328 | 21822401 | 21829520 | - | 13708 | 0.84 | 0.24 | 0.93 |
| chrUn_025064328 | 22182915 | 22187919 | + | 7222 | 0.79 | 0.31 | 0.91 |
| chrUn_025064328 | 22616744 | 22627212 | - | 25681 | 0.82 | 0.15 | 0.93 |
| chrUn_025064328 | 23480546 | 23487171 | - | 14788 | 0.83 | 0.38 | 0.92 |
| chrUn_025064328 | 24848015 | 24853713 | + | 9943 | 0.76 | 0.37 | 0.90 |
| chrUn_025064347 | 4711 | 11292 | + | 19264 | 0.87 | 0.29 | 0.91 |
| chrUn_025064389 | 3256519 | 3263718 | + | 16902 | 0.77 | 0.27 | 0.92 |
| chrUn_025064569 | 474985 | 480619 | . | 8077 | 0.88 | 0.37 | 0.93 |
| chrUn_025064569 | 518301 | 526775 | - | 16148 | 0.86 | 0.28 | 0.92 |
| chrUn_025064569 | 586919 | 596619 | - | 19150 | 0.83 | 0.27 | 0.86 |
| chrUn_025064569 | 623006 | 630719 | - | 17978 | 0.91 | 0.22 | 0.92 |
| chrUn_025064569 | 631712 | 640974 | - | 27551 | 0.90 | 0.22 | 0.92 |
| chrUn_025064569 | 691620 | 697110 | . | 8271 | 0.85 | 0.28 | 0.90 |
| chrUn_025064569 | 707805 | 712826 | - | 9765 | 0.88 | 0.28 | 0.94 |
| chrUn_025064569 | 736060 | 742494 | - | 21302 | 0.88 | 0.28 | 0.94 |
| chrUn_025064569 | 751914 | 763424 | - | 60312 | 0.91 | 0.31 | 0.93 |
| chrUn_025064569 | 846513 | 852303 | . | 12365 | 0.90 | 0.50 | 0.89 |

| | | | | | | | |
|---|---|---|---|---|---|---|---|
| chrUn_025064569 | 853629 | 866914 | - | 117939 | 0.92 | 0.30 | 0.91 |
| chrUn_025064569 | 870849 | 878823 | - | 74824 | 0.89 | 0.28 | 0.91 |
| chrUn_025064569 | 879435 | 1006512 | - | 1836023 | 0.90 | 0.25 | 0.92 |
| chrUn_025064569 | 1011837 | 1021318 | - | 34302 | 0.87 | 0.26 | 0.92 |
| chrUn_025064569 | 1021601 | 1039775 | - | 122068 | 0.87 | 0.31 | 0.93 |
| chrUn_025064569 | 2019339 | 2024510 | - | 6549 | 0.88 | 0.39 | 0.93 |
| chrUn_025064569 | 2438005 | 2445385 | + | 16752 | 0.76 | 0.27 | 0.92 |
| chrUn_025064719 | 1 | 5065 | - | 18498 | 0.89 | 0.19 | 0.94 |
| chrUn_025064737 | 5265869 | 5272074 | - | 10639 | 0.82 | 0.39 | 0.90 |

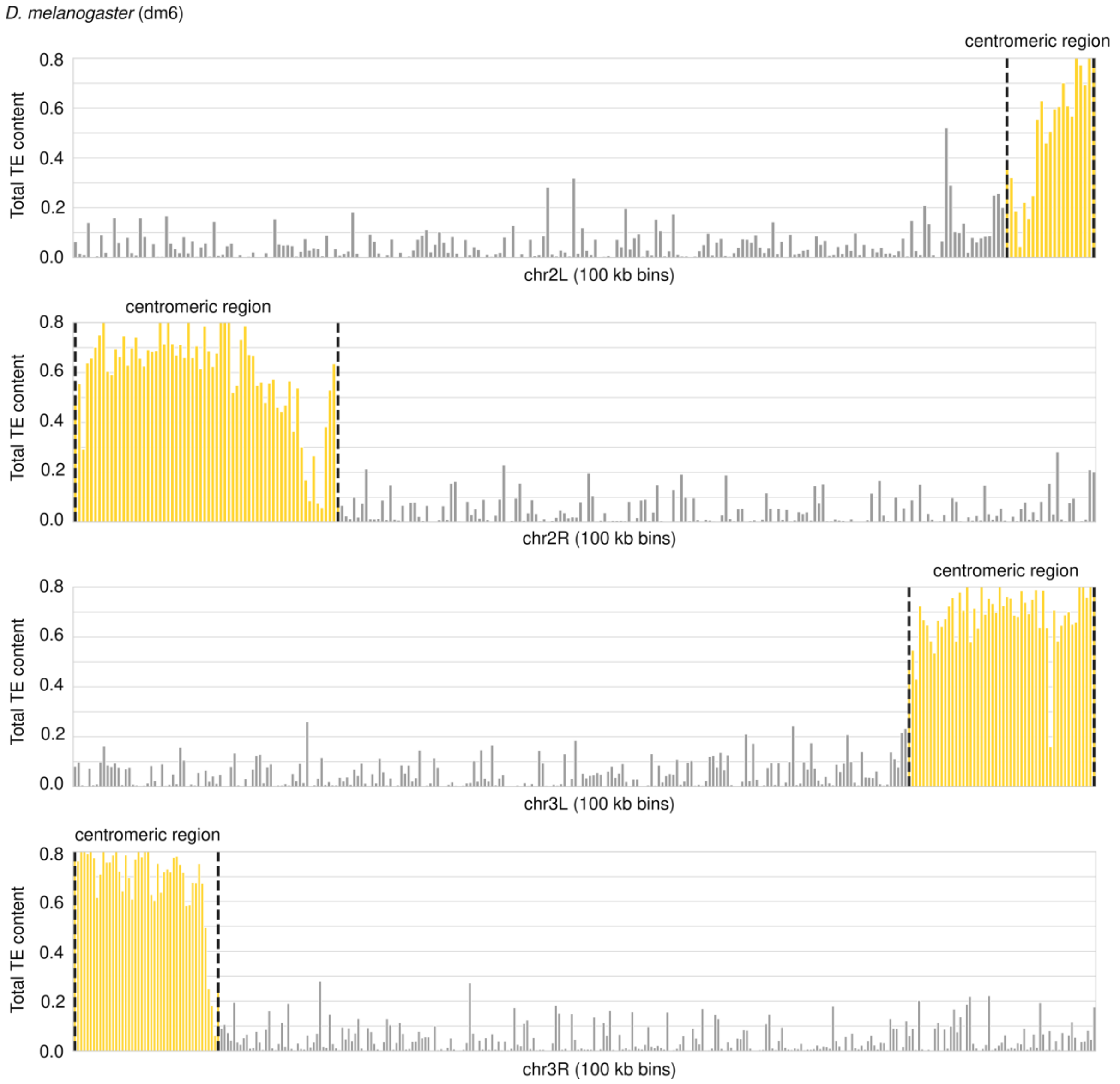


**Figure S1: Automatic annotation of centromeric regions.** Total TE content is shown across 100 kb bins on the left (L) and right (R) arm of chr2 and chr3 in *D. melanogaster*. A sliding window of five 100 kb bins was used to evaluate the TE content and centromeric regions were defined to start at the chromosome end and extend for as long as the TE content across the 5 bins was above 20 %. Centromeric regions are shown in yellow within the black dashed lines.

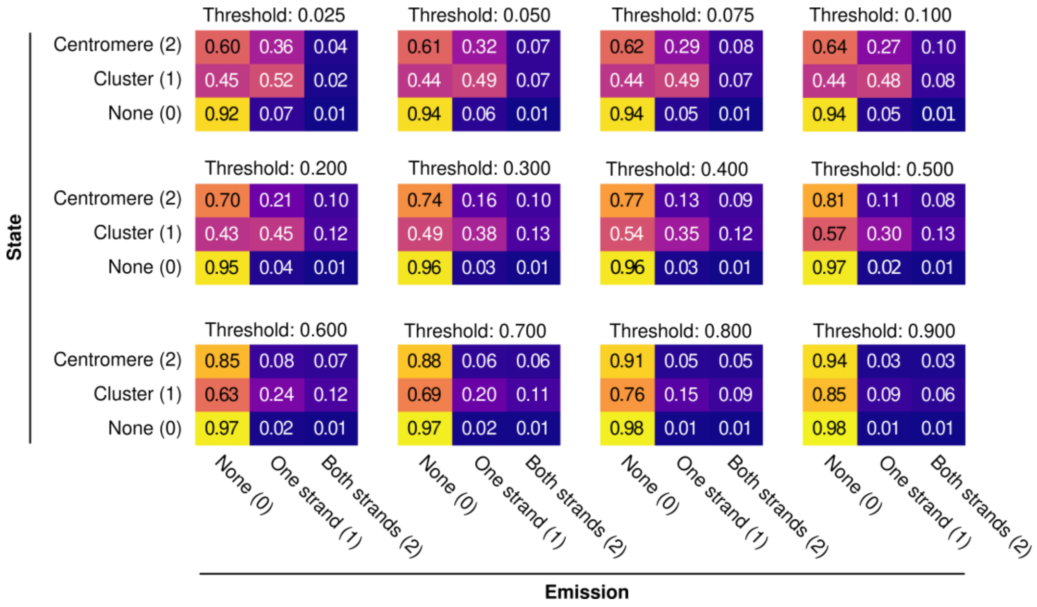


**Figure S2: Emission matrices depend on TE content threshold.** Emission matrices for *D. melanogaster* across different Gypsy content thresholds and using the 5 kb binning strategy. Emissions were defined based on the predicted LTR Gypsy content in each bin and categorised as: no Gypsy elements (None; 0), Gypsy elements present on one strand (One strand; 1), Gypsy elements on both strands (Both strands; 2).

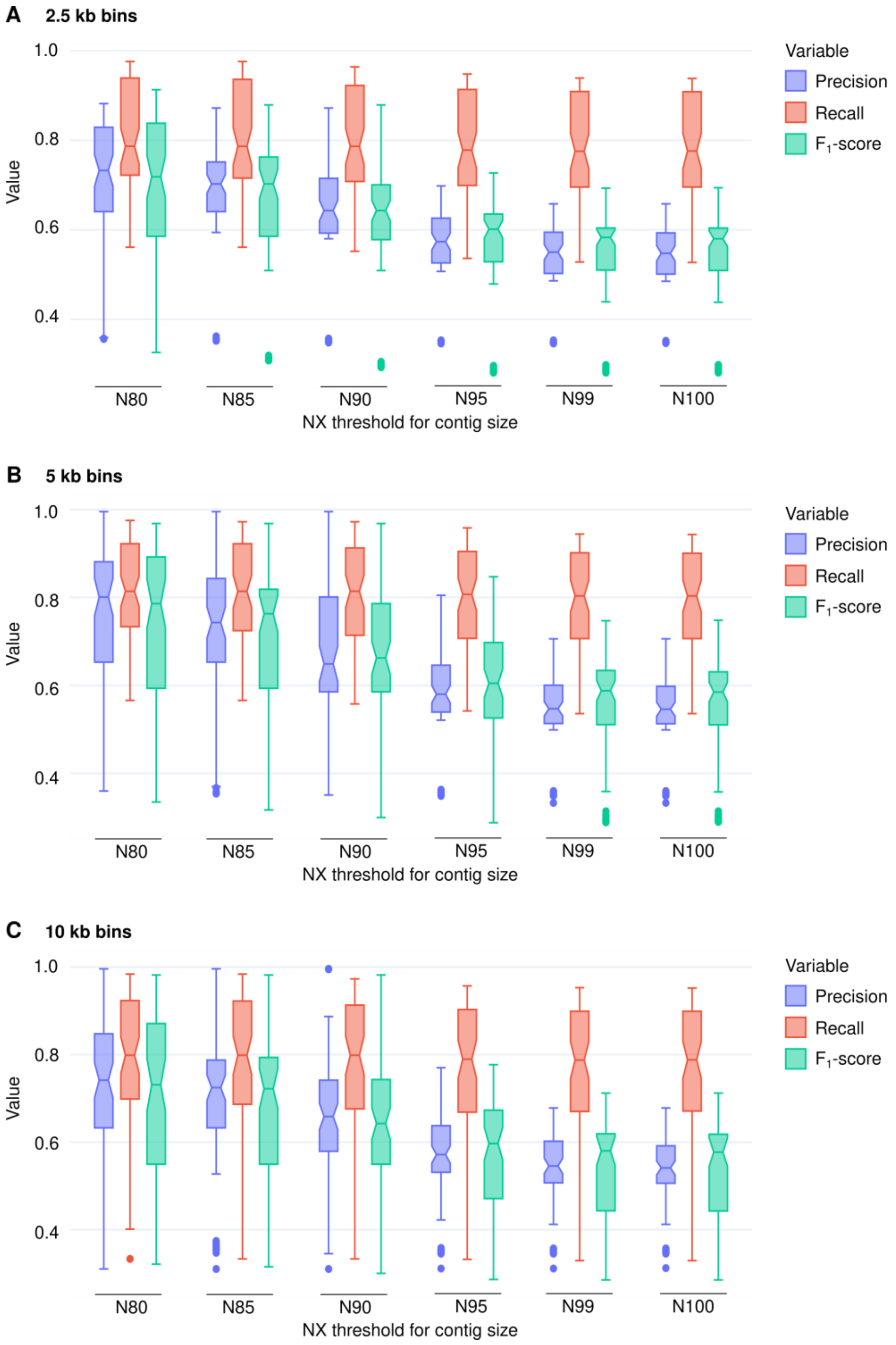


**Figure S3: Overview of model performance by NX threshold and bin size.** Cross validation model performance (mean across all three classes and all tested emission thresholds) using six different NX metric thresholds across (A) 2.5 kb, (B) 5 kb, or (C) 10 kb bins. Please note that N90 corresponds to the results presented throughout the study.

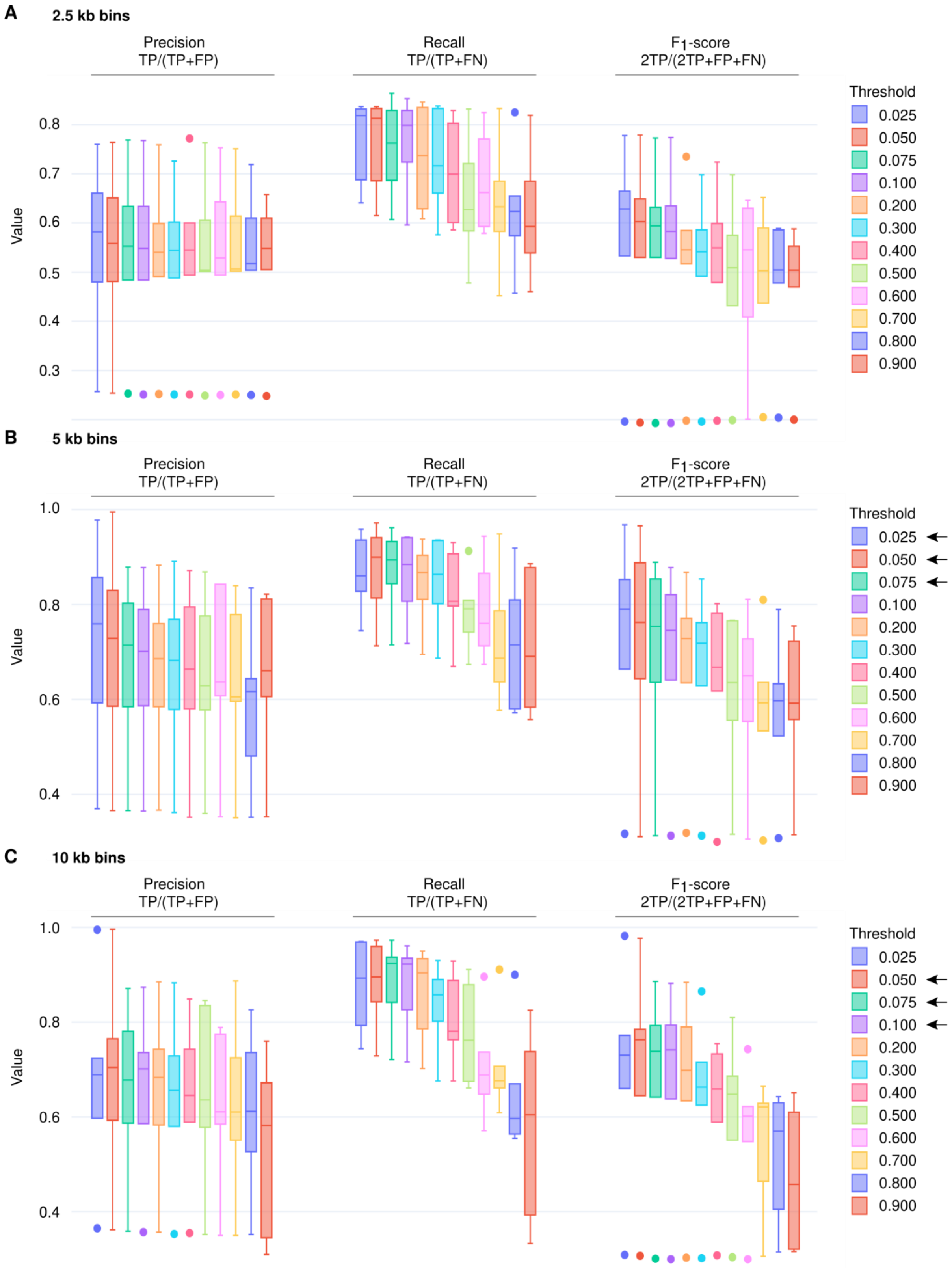


**Figure S4: Overview of model performance by threshold and bin size.** Cross validation model performance (mean across all three classes) using three evaluation metrics for 12 different thresholds for emission calculation across (A) 2.5 kb, (B) 5 kb, or (C) 10 kb bins. The thresholds with best $F_1$-scores are indicated with arrows.

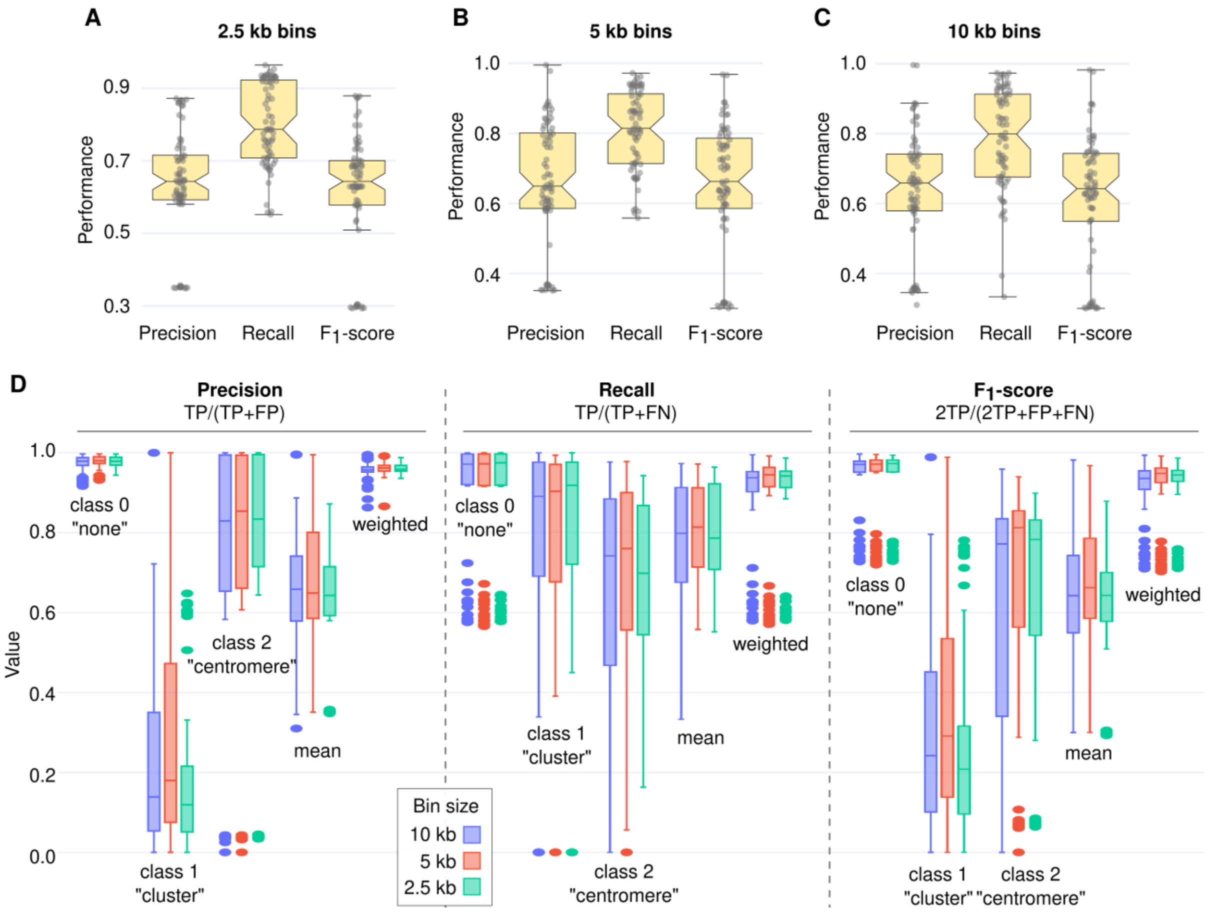


**Figure S5: Additional details on model performance.** Overall model performance based on cross validation using either (A) 10 kb or (B) 2.5 kb bins. (C) Detailed model performance with evaluation metrics for each class of the model (None, Cluster and Centromere), or the arithmetic or weighted mean using 10 kb, 5 kb, or 2.5 kb bins. All results are shown across all investigated emission thresholds.

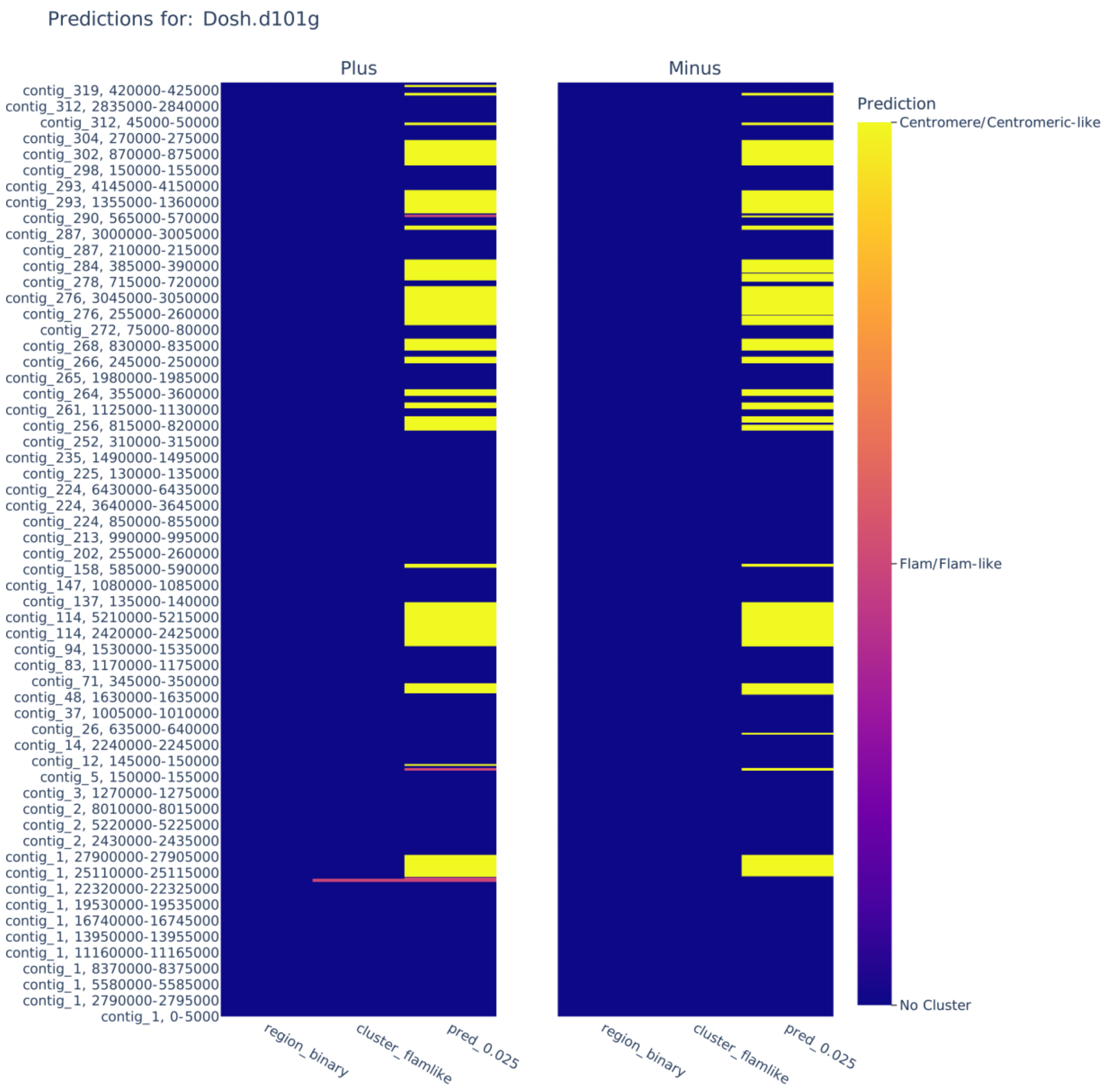


**Figure S6: Model output for *D. oshimai*.** Example of cluster prediction and visualisation for *D. oshimai* using bin size 5 kb and TE threshold 0.025. Annotations are shown as centromere (yellow), *flam*-like (pink), and none (blue) across N90 contigs.

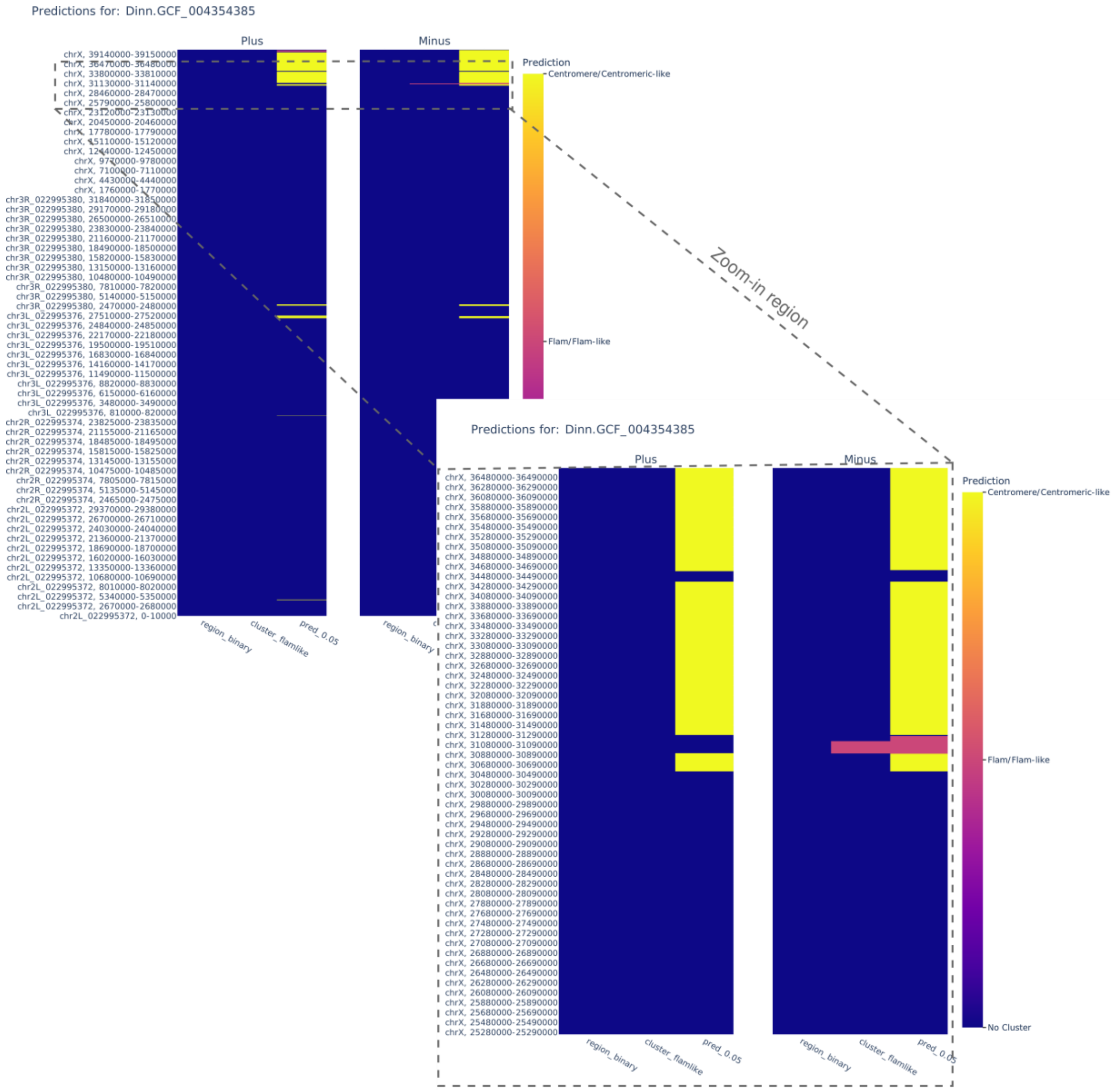


**Figure S7: Model output for *D. innubila*.** Example of cluster prediction and visualisation for *D. innubila* using bin size 10 kb and threshold 0.05. The visualisation allows the user to zoom interactively. An example zoom-in region is shown at the bottom right. Annotations are shown as centromere (yellow), *flam*-like (pink), and none (blue) across N90 contigs.

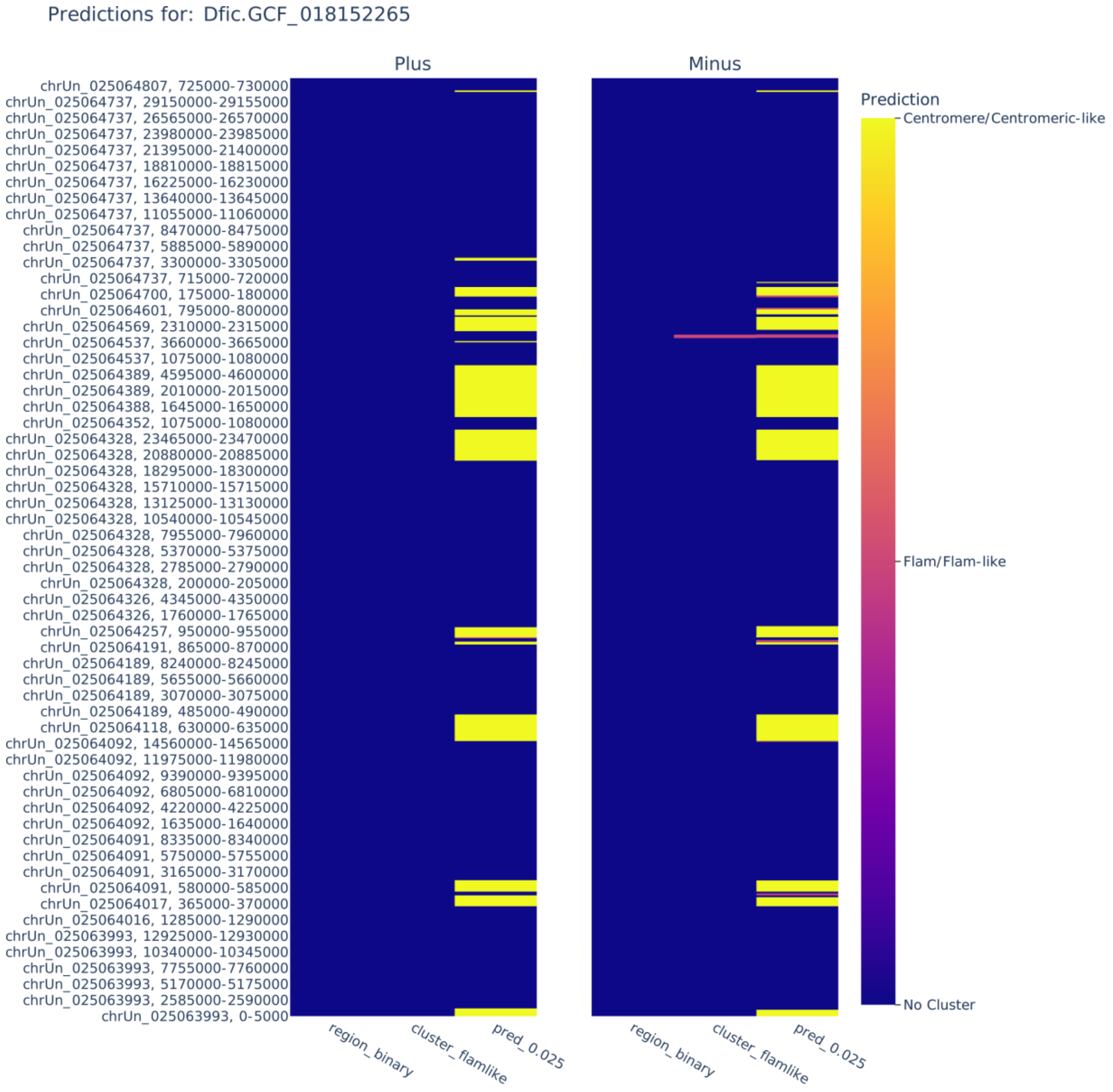


**Figure S8: Model output for *D. ficusphila*.** Example of cluster prediction and visualisation for *D. ficusphila* using bin size 5 kb and threshold 0.025. Annotations are shown as centromere (yellow), *flam*-like (pink), and none (blue) across N90 contigs.

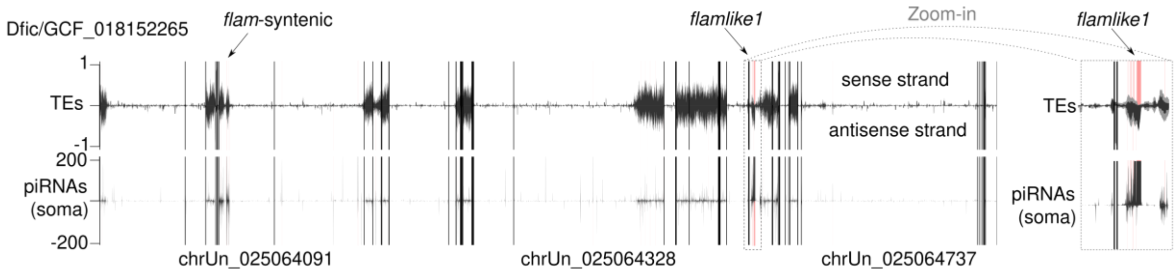


**Figure S9: Overview of proTRAC predictions for *D. ficusphila***. proTRAC piRNA cluster predictions across the *D. ficusphila* genome. Genomic coordinates are shown on the x axis with vertical black lines indicating contig breaks and selected contig names indicated. The top tracks show transposon content (black lines, LTR TEs; grey lines, all TEs) and the bottom one (black lines) shows piRNAs in soma-enriched ovarian cells (van Lopik et al., 2023). Positive values represent the sense strand and negative values the antisense strand relative to *flamlike1*. proTRAC predictions are shown as shaded pink areas. The major piRNA cluster predicted corresponds to previously reported *flamlike1* (van Lopik et al., 2023), indicated by an arrow and shown in the zoom-in to the right (grey dashed lines).
